# Broad SARS-CoV-2 Neutralization by Monoclonal and Bispecific Antibodies Derived from a Gamma-infected Individual

**DOI:** 10.1101/2022.10.14.512216

**Authors:** Denise Guerra, Tim Beaumont, Laura Radić, Gius Kerster, Karlijn van der Straten, Meng Yuan, Jonathan L. Torres, Wen-Hsin Lee, Hejun Liu, Meliawati Poniman, Ilja Bontjer, Judith A. Burger, Mathieu Claireaux, Tom G. Caniels, Jonne L. Snitselaar, Tom P. L. Bijl, Sabine Kruijer, Gabriel Ozorowski, David Gideonse, Kwinten Sliepen, Andrew B. Ward, Dirk Eggink, Godelieve J. de Bree, Ian A. Wilson, Rogier W. Sanders, Marit J. van Gils

## Abstract

The worldwide pandemic caused by SARS-CoV-2 has remained a human medical threat due to the continued evolution of multiple variants that acquire resistance to vaccines and prior infection. Therefore, it is imperative to discover monoclonal antibodies (mAbs) that neutralize a broad range of SARS-CoV-2 variants for therapeutic and prophylactic use. A stabilized autologous SARS-CoV-2 spike glycoprotein was used to enrich antigen-specific B cells from an individual with a primary Gamma variant infection. Five mAbs selected from those B cells showed considerable neutralizing potency against multiple variants of concern, with COVA309-35 being the most potent against the autologous virus, as well as against Omicron BA.1 and BA.2. When combining the COVA309 mAbs as cocktails or bispecific antibody formats, the breadth and potency was significantly improved against all tested variants. In addition, the mechanism of cross-neutralization of the COVA309 mAbs was elucidated by structural analysis. Altogether these data indicate that a Gamma-infected individual can develop broadly neutralizing antibodies.

## INTRODUCTION

More than two years since the beginning of the Coronavirus Disease 2019 (COVID-19) pandemic, over 619 million infections and 6 million deaths caused by Severe Acute Respiratory Syndrome Coronavirus 2 (SARS-CoV-2) have been reported to WHO, resulting in massive effects on global health and society in general^1^. Fortunately, the accelerated development of multiple, safe and effective COVID-19 vaccines has contributed to prevention of severe disease, reduced viral spread, and reduced stress on healthcare systems^2^. Although vaccines remain the best approach to mitigate COVID-19, alternatives are needed for individuals at risk, such as the elderly and the immune-compromised, who may not develop an immune response strong enough to forestall serious disease outcomes.

One promising alternative is the prophylactic and therapeutic use of neutralizing monoclonal antibodies (mAbs). Multiple SARS-CoV-2-directed mAb products have been developed and authorized for the emergency treatment of mild-to-moderate COVID-19 cases. These comprise cocktail therapies, including casirivimab (REGN10933) in combination with imdevimab (REGN10987) by Regeneron Pharmaceutical, bamlanivimab (LY-COV555) plus etesevimab (LY-CoV016) by Eli Lilly Company, tixagevimab (COV2-2196) plus cilgavimab (COV2-2130) by AstraZeneca, and two monotherapies, sotrovimab (VIR-7831) and bebtelovimab (LY-CoV1404/LY3853113) developed by Vir Biotechnology with GlaxoSmithKline and Eli Lilly, respectively^3^. The target of these therapeutics is the homotrimeric spike (S) glycoprotein exposed on the virion surface, which mediates the fusion of viral and host membranes after attachment to the human Angiotensin Converting Enzyme 2 (ACE-2) receptor^4, 5^. On the S protein, the major target of the neutralizing response is the Receptor Binding Domain (RBD), which transiently shifts between up and down conformations, thus representing a target where some epitopes are differentially exposed depending on RBD state^4^. Other regions of the S protein that are recognized by antibodies are the N-terminal domain (NTD) of the membrane-distal S1 head domain, and the conserved membrane-proximal S2 subunit that houses the fusion machinery and is generally targeted by more broadly reactive mAbs^6–9^.

More recently, the evolution of the wild-type (WT) virus into multiple variants that have acquired resistance to vaccines and mAb therapies has raised concerns about the longevity and efficacy of current treatment options, which are all based on the original Wuhan Hu-1 S sequence. Some of these variants are characterized by higher transmissibility, virulence and/or immune evasion compared to the WT virus, resulting in the WHO declaring the Alpha (Pango nomenclature B.1.1.7), Beta (B.1.351), Gamma (P.1), Delta (B.1.617.2) and Omicron (B.1.1.529) lineages as variants of concern (VOCs)^10^. In addition, variants of interest (VOIs) have also been reported worldwide. While the Alpha, Beta, Gamma and Delta variants have a moderate number of mutations in the S protein, the latest Omicron variant, with its sub-lineages BA.1, BA.2, BA.3, BA.4 and BA.5, contains more than 30 amino acid changes, of which almost half are located in the RBD, underpinning its substantial shift in antigenicity compared to early variants^11,12^. Several studies have indeed reported that the antigenic drift in the Omicron sub-lineages has resulted in a 20- to 40-fold reduction in the neutralization potency of sera from vaccinated and previously infected individuals^13–16^. In addition, a substantial or complete loss in neutralizing activity has also been observed for most commercial therapeutic mAbs, with sotrovimab, bebtelovimab and the combination of tixagevimab with cilgavimab being the only antibodies still showing activity against Omicron BA.1, albeit with a 5 to 100-fold loss of potency. Moreover, sotrovimab and tixagevimab plus cilgavimab have been shown to lose efficacy against Omicron BA.2 and BA.4/5^14–20^, and the emergence of resistance-associated mutations after sotrovimab administration have been recently reported^21,22^.

The marked reduction in VOCs neutralization by the current mAbs and vaccine-induced sera highlights the importance and the need for the discovery of novel broadly reactive neutralizing mAbs covering circulating and potential future emerging SARS-CoV-2 variants. In addition, although the isolation of mAbs from vaccinated and convalescent patients infected with the WT SARS-CoV-2 virus has been reported^6,23,24^, reports describing the discovery of such antibodies from VOC-infected individuals are limited^25^. Here, we describe the isolation of a set of human mAbs isolated from a convalescent unvaccinated COVID-19 patient who experienced a primary infection with the Gamma variant. The antibodies display a variety of functionalities against the autologous virus but also against heterologous VOCs, including WT, Alpha, Beta, Delta, Omicron BA.1 and BA.2. The specificities were corroborated by structural analyses. In particular, COVA309-35 had very potent neutralizing activity against Omicron BA.1 and BA.2 but less against Delta. Combining COVA309-35 with COVA1-18 and COVA1-16, previously isolated from a Wuhan Hu-1 infected individual^6^, yielded bispecific antibodies (bsAbs) with unusual breadth and potency.

## RESULTS

### Selection of SARS-CoV-2 spike-specific B cells and antibodies from a Gamma-infected individual

To study the B cell response induced by infection with the SARS-CoV-2 Gamma variant and isolate mAbs with novel specificities, we selected samples from an unvaccinated 32-year-old female, COSCA309, who experienced upper respiratory tract infection but did not require hospital admission, following a primary sequence-confirmed Gamma infection. COSCA309 was enrolled in the COSCA study (NL 73281.018.20) that was set up to facilitate investigation of VOC-specific B cell responses and isolation of broad and potent neutralizing mAbs against SARS-CoV-2 and its variants^6,11^.

We collected blood samples from COSCA309 approximately 40 days after symptom onset. After blood sample collection, peripheral blood mononuclear cells (PBMCs) were isolated and Gamma S-specific B cells were selected by flow cytometry-based single cell sorting using a stabilized Gamma S probe^6^ in two colors, in addition to the WT S protein in a third color. The flow cytometry analysis of the donor PBMCs showed a frequency of 0.58% Gamma S-specific B cells (Gamma S-AF647^+^, Gamma S-BV421^+^) among the total pool of B cells (CD19^+^Via^−^), which were shown to be predominantly memory B cells (CD27^+^IgD^−^, 61.5%) (Fig. 1a). Within these Gamma S-specific B cells, the majority expressed immunoglobulin G (IgG^+^) (57.3%), although a considerable portion of the S-specific B cells were IgM^+^ (34.7%). Co-staining with WT SARS-CoV-2 S protein indicated that 57.7% of Gamma S-specific B cells cross-reacted with the WT strain (Fig. 1a). After gene amplification and single-cell cloning, we obtained a total of 45 productive IgG heavy and light chain (HC/LC) pairs. The genetic signatures of the Gamma S-specific B cells were compared to the International Immunogenetics Information System (IMGT) germline repertoire^26^ (Supplementary Table 1). An overall increase in the usage of certain genes, such as IGHV3-53/3-66, IGHV1-2, IGHV3-30 and IGHV1-69, has been previously reported in COVID-19 patients infected with the WT SARS-CoV-2 virus^6,27–34^. Strikingly, none of the mAbs isolated in this study has been found to use the most dominant IGHV3-53 gene segment (Supplementary Table 1), whereas IGHV1-69 was found to be frequently used by the sorted Gamma S-specific B cells (15.6%), followed by IGHV3-7 and IGHV3-23 (11.1% each), and IGHV4-59 (8.9%) (Fig. S1a). The median somatic hypermutations (SHM) was 2%, in line with SHM levels observed after infection with the WT SARS-CoV-2^6,24,35,36^. We did not find any substantial difference in CDRH3 length of isolated Gamma S-specific B cells (mean 16 amino acids) compared to the average length of 15 amino acids generally present in the human naïve repertoire^37^. Following expression of the 45 mAbs in human embryonic kidney (HEK)293T cells and subsequent screening of the supernatants by a flow cytometry-based binding assay against the autologous Gamma S, as well as against Beta and WT S, a total of 14 mAbs were selected based on binding potency and breadth for large scale protein production and purification (Fig. 1b).

**Figure 1.**
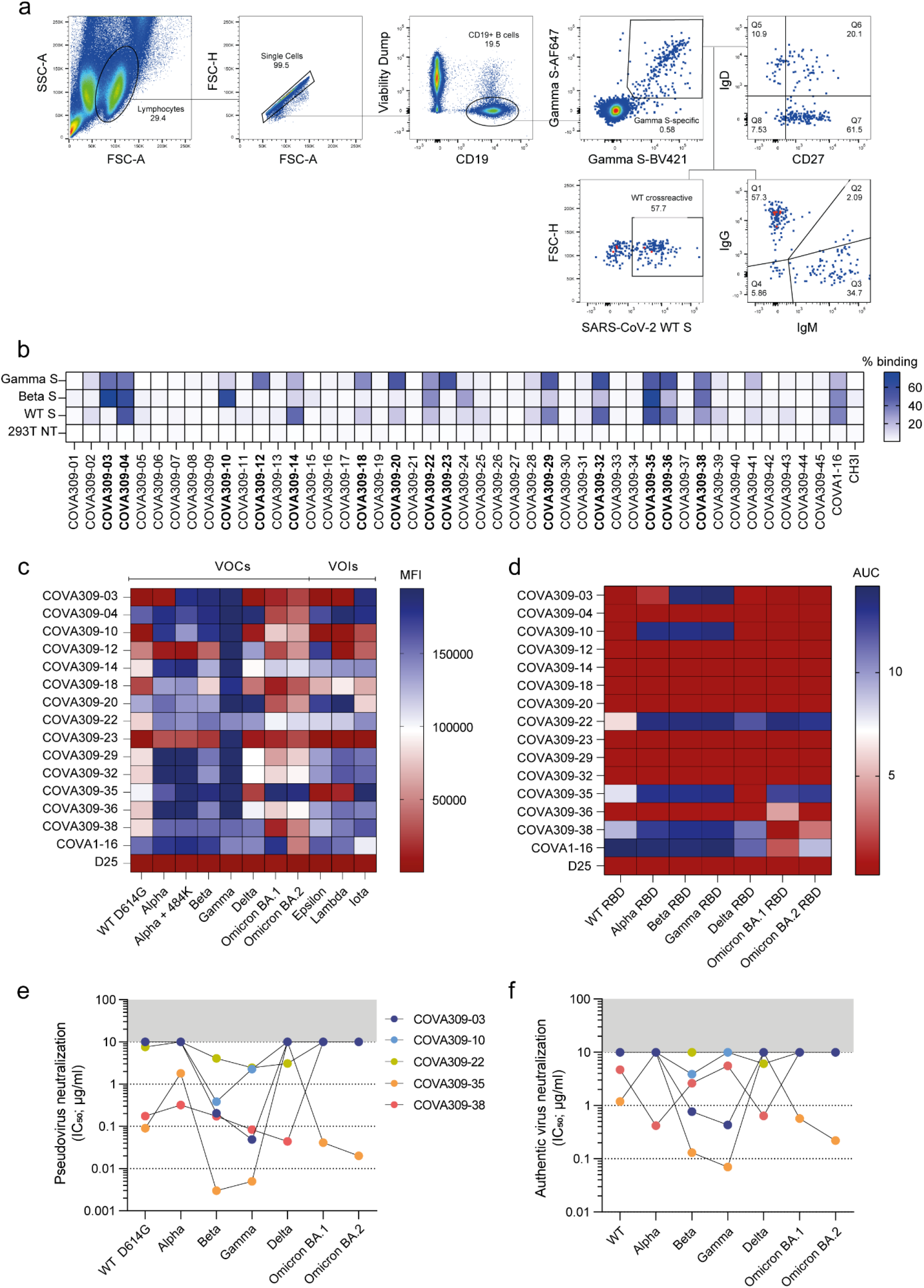
Selection of B cells and antibodies from a SARS-CoV-2 Gamma-infected individual. **a.** Sorting strategy of singlet viable CD19^+^ SARS-CoV-2 Gamma S-specific B cells. Gamma S-specific B cells were selected by double staining of Gamma S labelled with two different fluorescent dyes (Gamma S-AF647, Gamma S-BV421). In addition, Gamma S positive B cells were stained for IgD, CD27, IgG and IgM expression. Frequency of WT S cross-reactive B cells is also indicated. SSC-A, side scatter area; FSC-H, forward scatter height; FSC-A, forward scatter area. **b.** Flow cytometry-based screening of HEK293T-produced non-purified COVA309 supernatants against Gamma, Beta and WT S expressed on HEK293T cells, shown in the percentage of binding. 293T NT, non-transfected cells. COVA1-16 and CH3I antibodies were included as positive and negative controls, respectively. 14 mAbs (in bold) were selected for larger scale expression. **c.** Heat map showing the mean fluorescence intensity (MFI) of HEK293F-produced and purified COVA309 mAbs binding to SARS-CoV-2 variant S expressed on HEK293T cells, as assessed by flow cytometry. VOCs, variants of concern; VOIs, variants of interest. COVA1-16 and D25 (a RSV F specific mAb) are used as positive and negative controls, respectively. **d.** Heat map depicting the binding of COVA309 mAbs to SARS-CoV-2 variant RBDs, as determined by ELISA. Colour scale indicates the area under the curve (AUC) for each mAb. **e.** Half-maximal inhibitory concentrations (IC_50_) of SARS-CoV-2 VOC pseudoviruses neutralization for COVA309 mAbs. The cut-off was set at 10 μg/ml (grey bar). **f.** Neutralization of authentic SARS-CoV-2 viruses by COVA309 mAbs. The cut-off was set at 10 μg/ml (grey bar). Colour code is the same as for the pseudovirus neutralization.

We first assessed COVA309 mAbs binding specificities. Using flow cytometry, we tested the 14 antibodies for binding to HEK293T-transfected cells expressing a large panel of full-length membrane-expressed S, including WT D614G, VOCs (Alpha, Alpha with E484K, Beta, Gamma, Delta, Omicron BA.1 and BA.2), and VOIs (Epsilon, Lambda and Iota) (Fig. 1c). Beside binding to the autologous Gamma S protein, 11 out of the 14 mAbs also recognized the Beta variant, in line with the presence of high similarity between the S mutations in these strains. In general, COVA309 mAbs showed reduced binding to the WT D614G S compared to Beta and Gamma variants, while most of them still recognized the Alpha S. The majority of the mAbs showed decreased binding to Delta, Omicron BA.1 and BA.2 variants. COVA309-35 was unique since it showed strong binding to the Omicron BA.1 and BA.2 sub-lineages. In total, 7 mAbs recognized Epsilon, Lambda and Iota VOIs. In addition to the flow cytometry binding data to full-length S, we performed an enzyme-linked immunosorbent assay (ELISA) with soluble RBD proteins of the WT, Alpha, Beta, Gamma, Delta, Omicron BA.1 and BA.2 variants, and showed that 5 COVA309 mAbs are directed towards the RBD, while the others probably target other epitopes on the S, such as the NTD and the S2 subunit (Fig. 1d). Among the RBD-targeting mAbs, COVA309-22, −35 and −38 were the broadest, while COVA309-03 and −10 seemed to be more strain-specific. Overall, these data indicate that following a primary infection with the Gamma variant, broadly reactive mAbs against other viral strains can be elicited.

### Gamma-elicited antibodies show potent and broad neutralization

In addition to the binding data, neutralizing activity of the 14 selected mAbs was assessed in a lentiviral-based pseudovirus neutralization assay against the autologous Gamma strain (Fig. S1b). The RBD-targeting COVA309-03, −35 and −38 mAbs exhibited considerable neutralizing potency against the autologous Gamma variant, with half maximal inhibitory concentrations (IC_50_) ranging from 5 ng/ml to 84 ng/ml (Fig. 1e). Among them, COVA309-35 was the most potent (IC_50_ 5 ng/ml). In addition, two other mAbs (COVA309-10 and −22) were able to neutralize the autologous strain (IC_50_ 2.28 μg/ml and 2.45 μg/ml, respectively) (Fig. 1e). Since we aimed at finding potent cross-neutralizing mAbs, we tested the five COVA309 mAbs for broad neutralization against heterologous SARS-CoV-2 lineages, including the WT D614G, Alpha, Beta, Delta, Omicron BA.1 and BA.2 strains (Fig. 1e). Compared to autologous Gamma neutralization, we observed similar potencies against Beta, suggesting a primary role of key amino acid mutations shared between these two lineages, mainly located in the RBD (K417N/T, E484K and N501Y). Overall, we observed a marked reduction in Delta, Omicron BA.1 and BA.2 neutralization for the five COVA309 mAbs, consistent with the binding data. However, COVA309-35 still retained potent neutralization activity against the Omicron BA.1 and BA.2 variants (IC_50_ 41 ng/ml and 20 ng/ml, respectively), while it was less effective against Alpha (IC_50_ 1.8 μg/ml) and did not neutralize the Delta strain. On the contrary, antibody COVA309-38 strongly neutralized the Delta variant (IC_50_ 44 ng/ml), but not Omicron BA.1 and BA.2.

Next, we assessed the ability of the five COVA309 mAbs to neutralize authentic SARS-CoV-2 viruses, including WT, Alpha, Beta, Gamma, Delta, Omicron BA.1 and BA.2 strains (Fig. 1f). COVA309-03 neutralized primary Beta and Gamma viruses with an IC_50_ of 0.77 and 0.43 μg/ml, respectively, whereas COVA309-10 only neutralized Beta (IC_50_ 3.89 μg/ml). COVA309-35 and −38 were still highly effective in neutralizing multiple primary viruses. COVA309-35 neutralized autologous Gamma virus efficiently (IC_50_ 0.07 μg/ml) but showed a decrease in neutralization against Alpha and Delta viruses (IC_50_ >10 μg/ml), while maintaining neutralization of Omicron BA.1 and BA.2 variants (IC_50_ 0.57 and 0.22 μg/ml, respectively). COVA309-38 neutralized primary variants with IC_50_ ranging from 0.42 μg/ml to 5.54 μg/ml but was not effective against the Omicron BA.1 and BA.2 strains (IC_50_ >10 μg/ml). Compared to the pseudovirus neutralization, the neutralization potency against authentic SARS-CoV-2 strains was generally reduced, consistent with previous studies^6,38^, but the five neutralizing mAbs were overall similar between primary virus and pseudovirus neutralization assays in terms of activity and patterns of neutralization.

When we examined the genetic signatures of the five neutralizing versus non-neutralizing COVA309 mAbs, we found that COVA309-03, −10, −22 and −38 used genes belonging to the IGHV3 family (IGHV3-48, IGHV3-23, IGHV3-33 and IGHV3-7, respectively), while COVA309-35 was encoded by the dominant IGHV1-69 gene. The five mAbs used diverse IGKV/IGLV genes (IGKV1-5 for COVA309-03, IGKV1-39 for COVA309-10 and −22, IGKV1-27 for COVA309-35 and IGLV6-57 for COVA309-38). The neutralizing mAbs had low levels of SHM (median 1.7 %) and average heavy and light CDR3 lengths (10-23 amino acids and 9-10, respectively). These properties were similar to those of the Gamma-elicited non-neutralizing mAbs and previously described RBD-targeting mAbs that also present low SHM. However, the genetic features of mAbs isolated after WT infection, which predominantly use gene segments IGHV3-53/3-66 and IGHV3-30^6,35,37^, were found to differ from the gene usage of Gamma-elicited mAbs (Fig. S1a, Supplementary Table 1). Together these findings indicate that, despite certain features being preserved between WT- and VOC-induced mAbs, other signatures seem to be strain-specific.

### Structural analysis of COVA309 mAbs explains their ability to cross-neutralize

To study the mechanism of binding of the neutralizing COVA309 mAbs, we performed a biolayer interferometry (BLI) assay measuring competition between COVA309 mAbs and recombinant soluble human ACE-2 receptor for binding to autologous Gamma S (Fig. 2a). COVA309-03, −10 and −35 were able to block ACE-2 binding to Gamma S, suggesting that they target an epitope in close vicinity to the ACE-2-binding site on the apical area of the RBD. Alternatively, depending on the angle of approach to the RBD, they may cause steric hindrance with the receptor, therefore precluding ACE-2 interaction with S. These data are consistent with observations that many potent RBD-specific mAbs neutralize the virus by binding to a region that mediates ACE-2 receptor engagement, or by inhibiting the receptor attachment through specific approaching angles^39,40^. On the contrary, mAbs COVA309-22 and −38 were unable to block ACE-2 binding, suggesting that they might have different binding sites and mechanisms of action than ACE-2 blocking (Fig. 2a, 2c).

**Figure 2.**
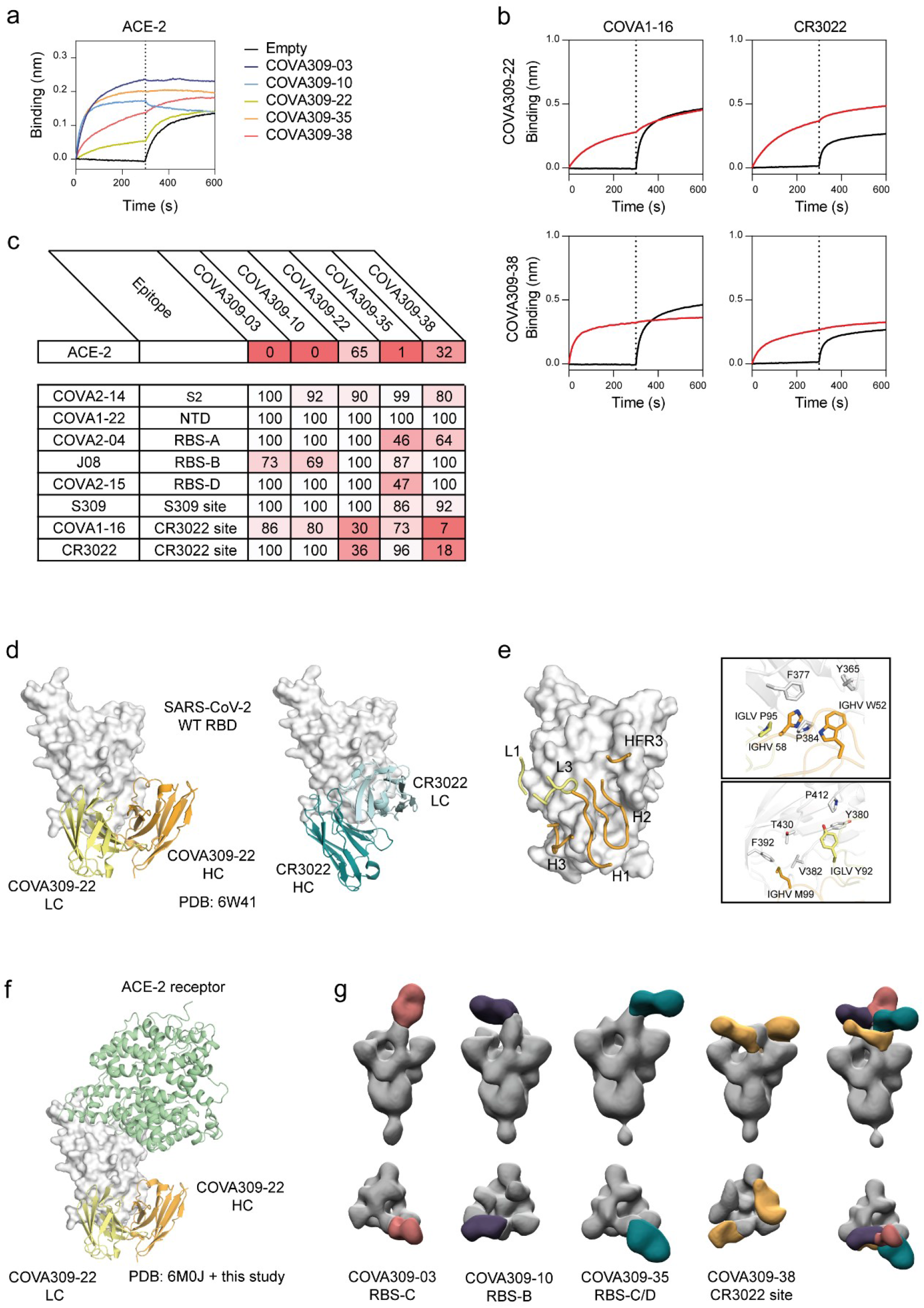
COVA309 mAb epitope determination. **a.** Biolayer interferometry plot depicts competition between the COVA309 mAbs and ACE-2, for binding to Gamma S coated on the chip. The black curves represent the baseline. **b.** Example of BLI plots showing COVA309-22 and −38 mAbs competing with COVA1-16 and CR3022 for binding to the WT S. The black curves represent the baseline, when no analyte was added. **c.** Heat map showing the percentage of residual binding of the ACE-2 receptor and other known SARS-CoV-2 mAbs after competition with COVA309 mAbs, as determined by BLI assay. For the mAbs competition, WT S was loaded on biosensors, and COVA2-14 (S2 binder), COVA1-22 (NTD binder), COVA2-04 (RBS-A), J08 (RBS-B), COVA2-15 (RBS-D), S309 (S309 site), COVA1-16 (CR3022 site) and CR3022 (CR3022 site)^6,41–43^ were included as competitors. **d.** Crystal structure of COVA309-22 Fab in complex with SARS-CoV-2 WT RBD at a 3.7-Å resolution (left). The HC and LC are coloured in orange and yellow, respectively. CR3022 Fab (right) is reported as a comparison (PDB: 6W41). **e.** Detailed representation of the main residues involved in the COVA309-22-WT RBD interaction. **f.** COVA309-22 HC (orange) and LC (yellow) bind the base and lateral face of the RBD, far from the ACE-2 receptor binding site (green; PDB: 6M0J). **g.** Front view (top row) and top view (bottom row) of low-resolution NS-EM reconstructions of COVA309-03, −10, −35 and −38 in complex with either Omicron 6P S or Gamma 6P S. Composite NS-EM maps of COVA309 Fabs indicate that all COVA309 mAbs align on one RBD in the up conformation.

Next, to acquire more insight into the binding site specificities of the COVA309 mAbs, we tested their binding by BLI to the WT SARS-CoV-2 S coated on biosensors and determined cross-competition with previously characterized mAbs. The mAbs directed to the RBD can be classified based on their binding epitope, targeting either sub-sites in receptor binding site, (RBS)-A, −B, −C, −D, or the more conserved S309 and CR3022 sites^41,42^. We included mAbs belonging to distinct classes, comprising COVA2-14 against S2, COVA1-22 against NTD, COVA2-04 against RBS-A, J08 against RBS-B, COVA2-15 against RBS-D, S309 against the S309 site, COVA-16 and CR3022 against the CR3022 epitope^6,41–43^ (Fig. 2b-c, S2a). COVA309-22 and −38 strongly interfered with binding of COVA1-16 and CR3022, indicating that they target the CR3022 site on the lateral face of the RBD, which is highly conserved among variants (Fig. 2b-c). COVA309-35 competed with COVA2-04 and COVA2-15 (46% and 47% residual binding, respectively), suggesting that its epitope partially overlaps with the regions targeted by the mentioned mAbs, or that the binding of COVA309-35 causes steric hindrance or induces a conformation change of the S which reduces the binding of COVA2-04 and COVA2-15. While we did not observe substantial competition of COVA309-03 with any of the tested mAbs, COVA309-10 showed a partial competition with RBS-B-targeting mAb J08.

We next determined the structural features and antigen recognition of the isolated COVA309 mAbs. We only obtained a crystal structure of the COVA309-22 antibody binding fragment (Fab) in complex with SARS-CoV-2 WT RBD at 3.7-Å resolution. In agreement with the BLI data, COVA309-22 interacts with a conserved RBD site that largely overlaps with that of CR3022 but in a different binding mode^42^ (Fig. 2d). The HC and LC of COVA309-22 and CR3022 bind the RBD with a swapped orientation, suggesting that, although the footprint is the same, the residues involved in the antigen recognition and interaction are different. The three CDRs of the COVA309-22 HC (CDRH1, H2 and H3) account for the majority of the interactions, burying 31 Å^2^, 377 Å^2^ and 232 Å^2^ of surface area (BSA), respectively, while the CDRL1, CDRL3 and HFR3 contribute to 73 Å^2^, 280 Å^2^ and 64 Å^2^ of the total 1057 Å^2^ BSA. Aromatic residues of COVA309-22 IGHV W52, H58, and IGLV P95 stack with RBD residues Y365, F377, and P384, whereas IGLV Y92 contacts RBD-Y380 and P412. Moreover, IGHV M99 forms hydrophobic interactions with RBD-V382, F392, and T430 (Fig. 2e). In accordance with competition experiments, the structure confirmed that the ACE-2 receptor binding site is far from the epitope recognized by this antibody and that COVA309-22 does not interfere with receptor binding (Fig. 2f).

In parallel, we generated lower resolution negative-stain electron microscopy (NS-EM) reconstructions of the other COVA309 mAbs. COVA309-03, −10 and −38 were analysed in complex with 6P-stabilized Gamma S, while COVA309-35 was complexed with 6P-stabilized Omicron BA.1 S protein (Fig. 2g, Fig. S2b). By using reference models from other mAbs, including CV07-270, CV07-250, C110 and DH1047^44,45^, we could determine the epitope for each mAb (Fig. S2c). In accordance with BLI competition data, COVA309-35 Fab in complex with the Omicron 6P S trimer showed that this antibody binds the apical part of the open, up-state RBD, targeting a region which encompasses the RBS-C and D regions^46,47^ (Fig. 2g). This corroborates the competition of COVA309-35 with COVA2-15 that binds RBS-D. In addition, RBS-C harbours L452, mutated in the Delta strain, explaining why COVA309-35 does not neutralize Delta. This finding is consistent with the ACE-2 competition data and shows that COVA309-35 resembles the binding characteristics of mAb C110, which was also reported to target a similar epitope^48^ (Fig. S2c). The NS-EM maps obtained from COVA309-03, −10 and −38 Fabs complexed with the Gamma 6P trimer demonstrated that all the antibodies could be aligned on the RBD in the up conformation, although COVA309-38 was also found to recognize the RBD in a partial down state. COVA309-10 binds RBS-B, on the apical part of the RBD, which includes amino acids at positions 478, 484 and 452, in accordance with our ACE-2 competition results and explains the loss in neutralizing activity against Delta, Omicron BA.1 and BA.2 that have different amino acids at these positions compared to Gamma. COVA309-03 targets RBS-C which also includes residues 484 and 452, thereby explaining the narrow neutralizing activity of this antibody. Lastly, NS-EM structures revealed that COVA309-38 binds the CR3022 site, in agreement with the competition to COVA1-16 and CR3022 itself (Fig. 2b-c, 2g).

### COVA309 mAbs contribute to potent and broad SARS-CoV-2 neutralization when incorporated into bispecific antibodies

Next, we investigated whether antibodies with complementary SARS-CoV-2 neutralizing potencies could be combined in a bispecific antibody (bsAb) format to achieve greater neutralization breadth. We focused on bsAbs that target distinct epitopes, including the most potent COVA309 mAbs, COVA309-35 and −38, and COVA1-18 and COVA1-16, two neutralizing mAbs isolated during the first wave of the pandemic^6,49,50^. We have previously reported on their distinct ability to recognize SARS-CoV-2 variants and indicated how COVA1-18 retains potency against Delta, while COVA1-16 is broadly reactive but with limited potency against Omicron strains^13^.

We generated a total of five bsAbs by combining COVA309-35 and −38 together or in combination with either COVA1-18 or COVA1-16, and studied the neutralizing potency of the bsAbs and corresponding antibody cocktails against WT D614G, Alpha, Beta, Gamma, Delta, Omicron BA.1 and BA.2 variants in a pseudovirus neutralization assay. Although COVA309-35 mAb alone remained the most potent antibody against the Beta and Gamma pseudoviruses (IC_50_ 3 and 5 ng/ml, respectively) (Fig. 1e), its ability to cross-neutralize other VOCs was significantly improved when combined in the bsAb format with COVA1-18, COVA1-16 or COVA309-38, gaining potency against Delta (IC_50_ 0.12, 2.18 and 0.99 μg/ml, respectively) and retaining strong neutralizing activity against Omicron BA.1 and BA.2, with IC_50_ values ranging from 0.05 to 2.7 μg/ml (Fig. 3a). Moreover, COVA309-35 together with COVA1-18, COVA1-16 and COVA309-38 neutralized WT D614G (IC_50_ 0.01, 0.13 and 0.09 μg/ml, respectively) and Alpha (IC_50_ 0.02, 0.33 and 0.13 μg/ml, respectively). When we tested bsAb COVA309-38 with COVA1-18 and COVA1-16, neutralization of all variants was improved compared to the parental COVA309-38 mAb (IC_50_ from 0.01 to 2.7 μg/ml), although activity against Omicron BA.1 and BA.2 was lower compared to bsAbs involving COVA309-35. For some combinations, in particular COVA309-35 together with COVA309-38 against Omicron BA.1 and BA.2, the corresponding antibody cocktails appeared superior compared to the bsAb formats (0.06 versus 2.7 μg/ml and 0.04 versus 0.15 μg/ml for cocktails and bsAbs against Omicron BA.1 and BA.2, respectively), indicating that antibody avidity may be important for the neutralization potency and suggesting that both Fab regions are needed for efficient virus neutralization and clearance, as previously shown for other mAbs^49,50^. Overall, these data indicate that when combined in multispecific formats, breadth of COVA309-35 and COVA309-38 are improved over the parental mAbs.

**Figure 3.**
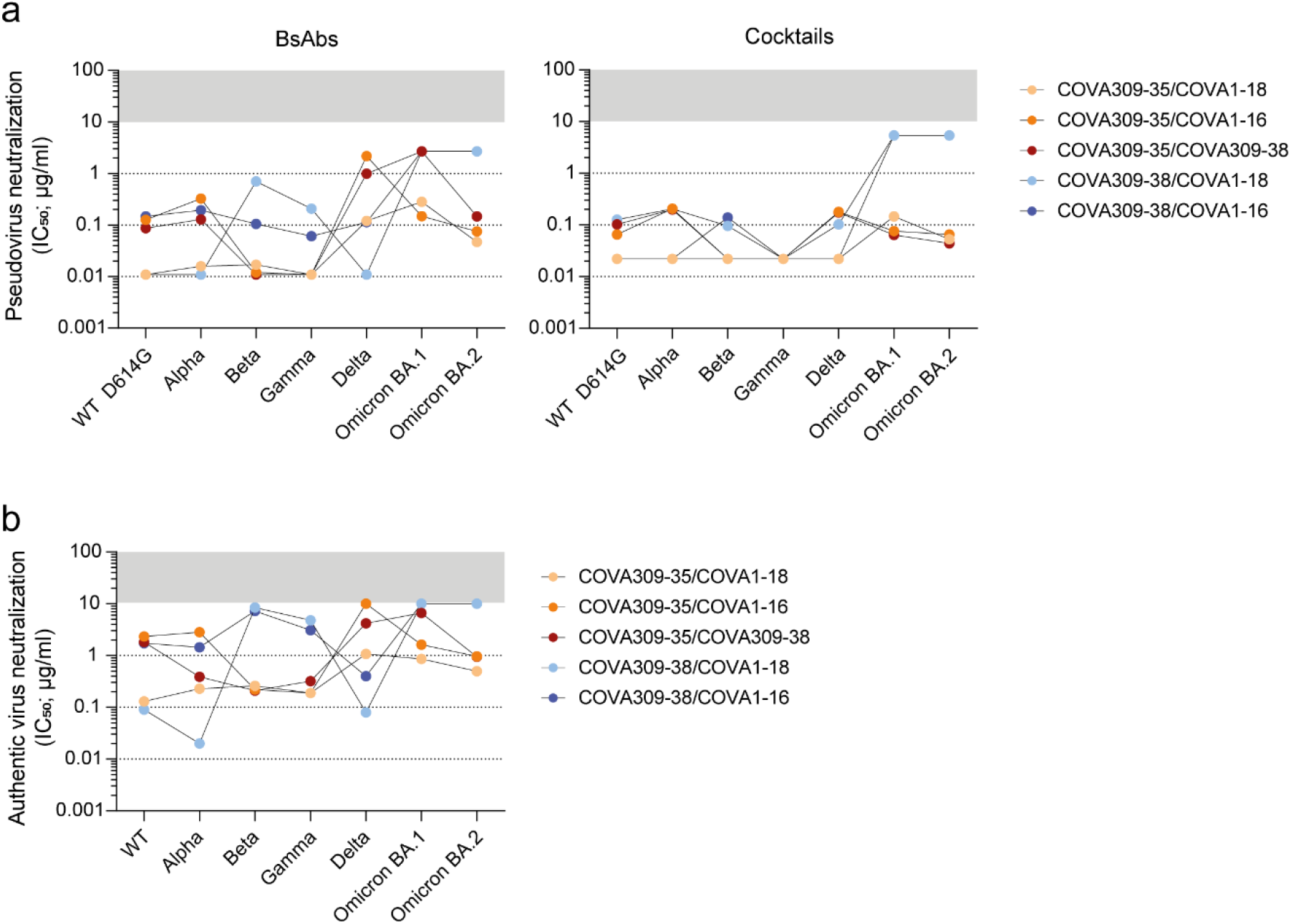
Characterization of bsAbs and antibody cocktails. **a.** IC_50_ values for pseudovirus neutralization of SARS-CoV-2 VOCs by bsAbs (left) and corresponding antibody cocktails (right). The cut-off was set at 10 μg/ml (grey bar). **b.** Neutralization of authentic SARS-CoV-2 WT, Alpha, Beta, Gamma, Delta, Omicron BA.1 and BA.2 viruses by bsAbs. The cut-off was set at 10 μg/ml (grey bar).

In addition to pseudovirus neutralization data, we also examined the neutralizing activity of the bsAbs against primary, authentic SARS-CoV-2 viruses, including WT, Alpha, Beta, Gamma, Delta, Omicron BA.1 and BA.2 (Fig. 3b). COVA309-35 in combination with either COVA1-18 or COVA309-38 appeared to have the highest potency and breadth, showing neutralization of all primary SARS-CoV-2 strains, including Omicron BA.1 and BA.2. COVA309-35 together with COVA1-16 also neutralized all variants except Delta, with a slightly lower potency (IC_50_ from 0.19 to 2.82 μg/ml). In accordance with the lack of neutralization capacity of COVA309-38 against Omicron BA.1 and BA.2, we did not observe any activity of the bsAbs involving COVA309-38 against these strains.

## DISCUSSION

SARS-CoV-2 has been shown to rapidly evolve into antigenically distinct variants, reducing the protective effect of natural infection as well as vaccination. The currently circulating SARS-CoV-2 Omicron variants that outcompeted the Delta lineage indeed show higher resistance to sera from naturally infected and/or vaccinated individuals^13–18^. Therefore, it is important to have potent therapeutics available, especially for the elderly and the immune-compromised. Different therapeutics have seen market introduction including several cocktails and mAbs. The mAbs, mAb cocktails or bsAbs should ideally be able to prevent viral escape and be broad enough to recognize diverse future SARS-CoV-2 lineages. However, most commercially available mAb products have been shown to lose potency against the latest Omicron sub variants^15,17–20^. Since we mainly depend on memory B cells for protection against serious disease and future outbreaks, it is important to study the immune repertoire of convalescent individuals to understand the breath and potency of the memory B cell response. In this study, we were able to demonstrate that Omicron and Delta neutralizing mAbs derived from memory B cells were present in a Gamma-infected individual. This finding is in contrast with the relatively poor cross-reactive serum response observed for COSCA309^11^, suggesting that the memory B cell response might be more versatile and therefore more adequately prepared to respond to future variants and be able to protect against severe disease.

The Gamma-elicited COVA309 mAbs differ substantially when comparing VOC binding, pseudovirus and primary virus neutralization. For example, COVA309-35 and −38 mAbs appear to have distinct S binding and neutralizing patterns against Omicron and Delta variants. Together, these mAbs can cover a broad range of antigenic variation within current and possible future SARS-CoV-2 strains. Therefore, we tested COVA309-35 and −38 together with COVA1-18 and COVA1-16 mAbs^6^, either as bsAbs or as cocktails in a pseudovirus neutralization assay (Fig. 3a). For bsAbs, lack of bivalent binding can be a disadvantage since the two different Fabs then should cooperate in binding to different epitopes on the same or adjacent S or S protomer. Although loss of bivalent binding can be observed for bsAbs compared to the combination therapy, the difference is small and, in general, both approaches provide broader antigenic coverage compared to the corresponding mAbs, especially the combination of COVA309-35 and COVA1-18. As a bsAb, this combination neutralizes all variants with IC_50_ between 0.01 and 0.28 μg/ml, with the latter value being for BA.1, while as a cocktail, the range is from 0.02 and 0.15 μg/ml, again the latter against BA.1.

In line with previous studies^47,48^, the binding epitopes of the five neutralizing COVA309 mAbs are mainly located in the immunodominant RBD of the S (Fig. 2c-g). More specifically, we show that our SARS-CoV-2 mAbs target distinct RBD sites, within and outside the RBS. The differences in neutralization of the VOCs by the COVA309 mAbs could be explained by the structural analysis and binding studies. COVA309-35, which showed the highest potency against the autologous Gamma strain, as well as against Omicron BA.1 and BA.2, interferes with ACE-2 receptor binding and recognizes an epitope covering RBS-C and D when the RBD is in the up state, thereby being highly effective against the infectious S conformation. COVA309-03 was also found to bind RBS-C, but its binding specificity appears to be narrower as it neutralizes only the Beta and Gamma variants. This indicates that its binding strongly depends on 484K, which was confirmed by recognition of an Alpha strain that we engineered to contain the E484K mutation (Fig. 1c).

In addition to mAbs targeting the RBS, COVA309-38 was found to target a more conserved cryptic site at the base and lateral face of the RBD. mAbs directed at this RBD region are generally broadly reactive but weakly neutralizing, although few exceptions exist like COVA1-16^6,49^ and ADI-62113^51^. COVA1-16 exhibits a special angle of approach and is able to directly compete with the receptor binding through steric hindrance, thereby neutralizing SARS-CoV-2 much more potently. As confirmed by BLI experiments and NS-EM, we showed that COVA309-38, in addition to COVA1-16, competes with CR3022 and COVA309-22 (Fig. 2b-c, Fig. S2d). In addition, ACE-2 competition data revealed that, despite targeting the base and lateral face of the RBD, this antibody is still able to partially compete with the receptor binding to the S, thereby in part resembling the mechanism of action of COVA1-16. The crystal structure of COVA309-22 showed that the antibody targets the same epitope as COVA309-38, explaining the breadth of the antibody, albeit with a lower potency compared to mAbs targeting the RBS^47^. The epitope of COVA309-22 also extensively overlaps with that of previously described mAbs, including CR3022^42^ and EY6A^52^, indicating once again that this RBD region represents a key site of vulnerability on the S protein.

Here, we demonstrate that potent and broadly reactive Gamma-elicited antibodies can be generated and can still neutralize the other variants, including the highly distant Omicron BA.1 and BA.2. We also show that, when tested in bsAb and cocktail combinations with other mAbs, all antibodies show enhanced neutralization breadth and potency, suggesting that combining antibodies with different RBD epitopes and mechanisms of action can provide a better resistance to viral mutants and therefore lead to more effective therapeutics. In addition, it is important to study how SARS-CoV-2 evolution shapes and alters the antibody response compared to the ancestral strain. In this way, we can increase our knowledge on the specific features and signatures of broadly reactive mAbs that cover diverse viral strains and may aid in development of next-generation vaccines and mAb therapeutics.

## ACKNOWLEDGMENTS

We thank the public health services (GGD) in the Netherlands for the help in contacting participants. We are also thankful to the participants of the COSCA study for the contribution to this research. We thank Robyn Stanfield for assistance in data collection. Use of the Stanford Synchrotron Radiation Lightsource, SLAC National Accelerator Laboratory, is supported by the U.S. Department of Energy, Office of Science, Office of Basic Energy Sciences under Contract No. DE-AC02-76SF00515. The SSRL Structural Molecular Biology Program is supported by the DOE Office of Biological and Environmental Research, and by the National Institutes of Health, National Institute of General Medical Sciences (P30GM133894).

## AUTHOR CONTRIBUTIONS

Conceptualization: R.W.S., M.J.vG., T.B., G.J.dB.

Funding acquisition: I.A.W., R.W.S., M.J.vG., G.J.dB.

Investigation: D.G., T.B., L.R., G.K., M.Y., J.L.T., W-H.L., H.L., M.P., I.B., J.A.B., J.L.S., S.K., D.G.

Methodology: L.R., K.S., M.C., T.G.C., D.E., G.O., A.B.W.

Resources: L.R., G.J.dB., K.vdS., D.E., T.G.C, I.B.

Supervision: G.O., A.B.W., T.B., K.S., M.C., I.A.W., R.W.S., M.J.vG.

Writing – original draft: D.G., T.B., R.W.S., M.J.vG.

Writing – review & editing: all authors

---

**Supplementary Table 1.** Genetic signatures of Gamma-elicited B cells. Table reporting the gene usage, V region identity (%) and CDRH3 sequence for the produced Gamma-specific B cells. VH and VL region identity refers to V gene segment identity compared to the germline repertoire (International Immunogenetics Information System)^26^.

## MATERIALS AND METHODS

### Patient sample

Blood from a SARS-CoV-2 Gamma-infected adult (COSCA309) was collected approximately 40 days after symptom onset through the COVID-19-specific antibodies (COSCA) study (NL 73281.018.20). The participant had a SARS-CoV-2 positive nasopharyngeal swab, as assessed by qRT-PCR (Roche LightCycler480, targeting the Envelope-gene 113bp). After density gradient centrifugation, peripheral blood mononuclear cells (PBMCs) of the donor were isolated. The COSCA study was conducted at the Amsterdam University Medical Centre, location AMC, the Netherlands and approved by the local ethical committee. Written informed consent was provided by the participant before being enrolled in the study.

### Protein constructs design and production

The mutations present in the stabilized S proteins compared to the WT strain (Wuhan Hu-1; GenBank: MN908947.3) are reported in Supplementary Table 3. Soluble RBD protein constructs of the different variants were designed with the same mutations reported in the table (residues 319-541). gBlock gene fragments (Integrated DNA Technologies) of the different constructs were ordered and cloned into a pPPI4 avidin-tagged and/or hexahistidine-tagged vector by PstI-BamHI digestion and ligation with Gibson Assembly (Thermo Fisher Scientific). The recombinant human ACE-2 receptor was obtained in the same way after ordering the corresponding gBlock gene fragment (Integrated DNA Technologies). Sanger sequencing was used to verify the constructs. Proteins were then produced in human embryonic kidney (HEK)293F cells (Thermo Fisher Scientific) maintained in Freestyle medium (Life Technologies). Briefly, HEK293F cells were transfected with a 1:3 ratio of expression DNA plasmids (312.5 μg/l) and Polyethylenimine Hydrochloride (PEI)max (1 μg/μl) in OptiMEM. Supernatants containing the produced proteins were harvested six days after transfection, centrifuged at 4000 rpm for 30 min and filtered using 0.22 μM Steritop filter units (Merck Millipore). Affinity chromatography with Ni-NTA agarose beads (Qiagen) was used to purify the proteins. Eluates were subsequently concentrated and buffer exchanged to phosphate-buffered saline (PBS) or TN75 buffer (75 mM NaCl and 20 mM Tris HCl, pH 8.0) using VivaSpin20 filters (Sartorius). Protein concentrations were determined by the Nanodrop.

### Probe staining and single B cell sorting

To produce fluorescent labelled-probes for the B cell sorting, soluble avidin-tagged Gamma S protein was biotinylated with a BIrA500 biotin-ligase reaction kit according to the manufacturer’s instruction (Avidity). Biotinylated protein was then mixed with streptavidin fluorophores (AF647, BioLegend; BV421, BioLegend), as described previously^6^, and incubated at 4°C for 1 h. 10mM biotin (Genecopoiea) was added for at least 10 min to quench unbound streptavidin conjugates. Labelled proteins were then mixed with PBMCs for 30 min at 4°C, washed with FACS buffer (PBS supplemented with 1 mM EDTA and 2% fetal calf serum) and stained with a live/dead marker (viability-eF780, eBiosciences) together with CD19-AF700 (HIB19, BioLegend), CD20-PE-CF594 (2H7, BD Biosciences), CD27-PE (L128, BD Biosciences), IgM-BV605 (MHM-88, BioLegend), IgG-PE-Cy7 (G18-145, BD Biosciences). Sample was washed twice and acquired on the ARIA-SORP-II 4 lasers for B cell sorting. The lymphocyte population was first gated based on the morphology (FSC-A and SSC-A) and doublets were removed. Live B cells (CD19^+^Via^−^) double positive for the SARS-CoV-2 Gamma S protein (AF647 and BV421) were single cell-sorted into 96-well plates containing lysis buffer (20 U RNAse inhibitor (Invitrogen), first strand SuperScript III buffer (Invitrogen), and 1.25 μl of 0.1 M DTT (Invitrogen)). After single cell sorting, the plates were stored at −80°C for at least 1 h before a reverse transcriptase (RT)-PCR was performed to convert the mRNA of the lysed B cells into cDNA, as described previously^6^. Analysis was performed using FlowJo X software (BD Biosciences).

### Single-cell immunoglobulin gene amplification, cloning and antibodies expression

After RT-PCR, additional PCR rounds were performed to amplify the V(D)J variable region of the HC and LC of the antibodies, as described previously^6^. PCR products were then cloned into corresponding human IgG1 expression vectors with Gibson Assembly (Thermo Fisher Scientific) and the mixture was subsequently transformed into DH5α cells. After DNA purification, the sequences were verified by Sanger sequencing. For small scale expression of the antibodies, adherent HEK293T cells (ATCC, CRL-11268) were cultured in Dulbecco’s Modified Eagle Medium (DMEM) supplemented with 10% foetal calf serum and a mixture of penicillin/streptomycin (100U/ml and 100 μg/ml, respectively). HEK293T cells were seeded in 24-well plates at a density of 2.75×10^5^ cells/well 24h prior to transfection. The transfection was performed with a 1:1 (w:w) HC/LC ratio using a 1:2.5 ratio with 1 mg/l PEImax (Polysciences) in 200 μl Opti-MEM. The transfection mix was incubated for 15 min at RT and then added onto the cells. Supernatants of transfected cells were harvested 48 h post-transfection and tested for preliminary screening by flow cytometry.

### Larger expression of antibodies in HEK293F cells

Suspension HEK293F cells (Invitrogen) were maintained in FreeStyle medium (Gibco) and co-transfected with a 1:3 ratio of the two HC/LC DNA plasmids and 1 mg/l PEImax (Polysciences). Five days post-transfection, the cell suspension was centrifuged at 4000 rpm for 30 min, followed by filtration of the supernatant using 0.22 μm pore size SteriTop filters (Millipore). The filtered supernatant containing the recombinant IgG antibodies was run over a 10 ml protein G column (Pierce) and the antibodies were then eluted with 0.1 M glycine pH 2.5, into the neutralization buffer (1 M TRIS pH 8.7). 50 kDa VivaSpin20 columns (Sartorius) were used to concentrate and buffer exchange the antibodies to PBS. The IgG concentration was determined by the NanoDrop.

### Flow cytometry-based screening of supernatants and purified recombinant mAbs

Cell surface-expressed SARS-CoV-2 ancestral and variant S were obtained by transfecting 8 μg of SARS-CoV-2 full-length DNA plasmid with 25 μl PEImax in 400 μl OptiMEM onto 12 to 15 ml HEK293T cells in a petri dish (seeded the day before at a density of 3.0×10^6^). After 48 h, cells were harvested and frozen until further use. After thawing, HEK293T cells expressing the S of interest were seeded at 20.000 to 30.000 cells per 96-well in FACS buffer (PBS/0.5% FCS) and were incubated 1:1 with non-purified supernatants from small scale IgG production or a dilution of purified HEK293F-produced mAbs for 30 min at 4°C. Subsequently, cells were washed twice with FACS buffer and incubated for 30 min at 4°C with 1:1000 diluted PE-conjugated goat F(ab)’2 anti-human IgG (Southern Biotech 2042-09) in the dark. Cells were washed and analysed on the FACS Canto II analyser. Samples were analysed by FlowJo X software (BD Biosciences) and percentage of cells that showed binding were plotted correspondingly.

### Pseudovirus design and neutralization assay

The WT D614G, Alpha, Beta, Gamma, Delta, Omicron BA.1 and BA.2 pseudovirus constructs were ordered as gBlock gene fragments (Integrated DNA Technologies) and cloned using Gibson Assembly (ThermoFisher), as described previously^6^. Briefly, all S constructs were produced by co-transfecting HEK293T cells with the plasmid expressing the S with the pHIV-1NL43 ΔEnv-NanoLuc reporter virus plasmid. Cell supernatant containing the pseudovirus was harvested 48 hours post-transfection and stored at −80°C until further use. To test the neutralization activity, mAbs, bsAbs and antibody cocktails were serially diluted in 3-fold steps starting at a concentration of 10, 2.7 and 5.4 μg/ml, respectively. They were then mixed with the pseudovirus and incubated for 1 h at 37°C. The mixes containing the pseudovirus and the mAbs/bsAbs were then added to HEK293T-ACE2 cells, seeded the day before in poly-L-lysine-coated 96-well plates at a density of 20.000 cells/well. After 48 h, the pseudovirus-mAbs/bsAbs combination was removed, cells were lysed and transferred to half-area 96-wells white microplates (Greiner Bio-One). Luciferase activity of cell lysate was measured using the Nano-Glo Luciferase Assay System (Promega) with a Glomax plate reader (Turner BioSystems).

### Authentic virus neutralization assay

We tested mAbs and bsAbs for their neutralization capacity against the ancestral SARS-CoV-2 virus (German isolate; GISAID ID EPI_ISL 406862; European Virus Archive Global #026V-03883) and VOCs, as previously described^53^. Briefly, samples were serially diluted in Dulbecco modified Eagle medium supplemented with NaHCO_3_, HEPES buffer, penicillin, streptomycin, and 1% fetal bovine serum, starting at a dilution of 10 μg/mL in 50 μl. Subsequently, 50 μL of virus suspension were added to each well and incubated at 35°C for 1 h. Vero E6 cells were added in a concentration of 20.000 cells per well and subsequently incubated for 48 hours at 35°C. After incubation, cells were fixed with 4% formaldehyde/phosphate-buffered saline (PBS) and stained with a nucleocapsid targeting monoclonal antibody. Bound ab as a measure for infected cells was detected using horseradish peroxidase–conjugated goat anti-human IgG (1:3000) in 2% milk/PBS for 1 hour at RT. After washing, the color reaction was developed using 3,3, 5,5’-tetramethylbenzidine substrate (Thermo Scientific Scientific). The reaction was stopped by adding 0.8 N sulfuric acid, and OD450 (optical density at 450 nm) was measured using standard equipment.

### Ni-NTA enzyme-linked immunosorbent assay

Soluble hexahistidine-tagged SARS-CoV-2 WT, Alpha, Beta, Gamma, Delta, Omicron BA.1, BA.2 RBD proteins (1μg/ml) were loaded in casein (Thermo Scientific) on 96-well Ni-NTA plates (Qiagen) for 2h at RT. After washing the plates with Tris-Buffered Saline (TBS), five-fold serial dilutions of mAbs (starting from 5 μg/ml) in casein were added to the plates. After incubating the plates for 2h at RT, they were washed three times with TBS, and a 1:3000 dilution of HRP-labelled goat anti-human IgG (Jackson Immunoresearch) in casein was added for 1h at RT. Plates were finally washed five times with TBS/20% Tween. Colorimetric detection was performed after developing the reaction for 4 min before termination by adding 0.8M sulfuric acid.

### Competition assays by biolayer interferometry

For competition experiments, an Octet K2 (ForteBio) was used. Briefly, to test the competition of the ACE-2 receptor with COVA309 mAbs, autologous hexahistidine-tagged Gamma S (10 μg/ml) was loaded in running buffer (PBS, 0.02% Tween20, 0.1% BSA) on Ni-NTA biosensors. To measure association, the chip was first dipped in running buffer to remove protein in excess, and subsequently dipped for 300s in a well containing 10 μg/ml mAb in running buffer. Next, the chip was dipped for 300s in 5 μg/ml of recombinant human ACE-2 in running buffer to measure competition with the mAbs.

For mAbs cross-competition, Ni-NTA biosensors were loaded with 10 μg/ml hexahistidine-tagged SARS-CoV-2 WT S protein in running buffer. After dipping the chip in running buffer to remove excess of S, the chip was transferred for 300 s to a well containing 10 μg/ml of one of the COVA309 mAbs in running buffer. Next, the chip was dipped for 300s in either 10 μg/ml of previously characterized COVA mAbs^6^ and CR3022^42^, or 5 μg/ml of J08 and S309 mAbs in running buffer to measure cross-competition.

### Bispecific antibodies production

BsAbs were generated by introducing amino acid mutations (F405L or K409R) in the HC constant region of the parental mAbs by Q5 Site-Directed Mutagenesis (New England Biolabs). Individual mutated mAbs were produced in HEK293F cells and purified as reported above. BsAb combinations were generated by using a previously described controlled Fab-arm exchange protocol^54^ and later tested for neutralization against SARS-CoV-2 VOC pseudo and primary viruses.

### Expression and purification of antibody fragments (Fabs)

The plasmids were transiently co-transfected into ExpiCHO cells at a ratio of 2:1 (HC:LC) using ExpiFectamine™ CHO Reagent (Thermo Fisher Scientific), according to the manufacturer’s instructions. The supernatant was collected at 10 days post-transfection. The Fabs were purified with a CaptureSelect™ CH1-XL Affinity Matrix (Thermo Fisher Scientific), followed by size exclusion chromatography.

### Expression and purification of SARS-CoV-2 RBD for crystallization

Expression and purification of the SARS-CoV-2 RBD for crystallization were performed as described previously^42^. Briefly, the RBD (residues 333-529) of the SARS-CoV-2 S protein (GenBank: QHD43416.1) was cloned into a customized pFastBac vector^55^ and fused with an N-terminal gp67 signal peptide and C-terminal His6 tag^42^. A recombinant bacmid DNA was generated using the Bac-to-Bac system (Life Technologies). Baculovirus was generated by transfecting purified bacmid DNA into Sf9 cells using FuGENE HD (Promega), and subsequently used to infect suspension cultures of High Five cells (Life Technologies) at an MOI of 5 to 10. Infected High Five cells were incubated at 28 °C with shaking at 110 rpm for 72 h for protein expression. The supernatant was then concentrated using a 10 kDa MW cutoff Centramate cassette (Pall Corporation). The RBD protein was purified by Ni-NTA, followed by size exclusion chromatography, and buffer exchanged into 20 mM Tris-HCl pH 7.4 and 150 mM NaCl.

### Crystallization and structural determination

The COVA309-22/RBD complex was formed by mixing each of the protein components at an equimolar ratio and incubating overnight at 4°C. The protein complex was adjusted to 11 mg/ml and screened for crystallization using the 384 conditions of the JCSG Core Suite (Qiagen) on our robotic CrystalMation system (Rigaku) at Scripps Research. Crystallization trials were set-up by the vapor diffusion method in sitting drops containing 0.1 μl of protein and 0.1 μl of reservoir solution. Optimized crystals were then grown in drops containing 0.1 M sodium cacodylate, pH 6.5, 0.2 M magnesium chloride, and 20% (w/v) polyethylene glycol 1000 at 20°C. Crystals appeared on day 3, were harvested on day 15 by soaking in reservoir solution supplemented with 20% (v/v) ethylene glycol, and then flash cooled and stored in liquid nitrogen until data collection. Diffraction data were collected at cryogenic temperature (100 K) at the Stanford Synchrotron Radiation Lightsource (SSRL) on Scripps/Stanford beamline 12-2. Diffraction data were processed with HKL2000^56^. The model of COVA309-22 was generated by Repertoire Builder^57^. Structures were solved by molecular replacement using PHASER^58^. Models for molecular replacement were derived from PBD 6W41. Iterative model building and refinement were carried out in COOT^59^ and PHENIX^60^, respectively. Epitope and paratope residues, as well as their interactions, were identified by accessing PISA at the European Bioinformatics Institute^61,62^.

### Negative stain electron microscopy analysis

SARS-CoV-2 S protein was complexed with three-fold molar excess of Fab and incubated at RT for 30 min. The complex was diluted to 0.02 mg/ml in 1xTBS and 3 μl applied to a 400mesh Cu grid, blotted with filter paper, and stained with 2% uranyl formate. Micrographs were collected on a ThermoFisher Tecnai Spirit microscope operating at 120kV with a FEI Eagle CCD (4k) camera at 52,000 magnification using Leginon automated image collection software^63^. Particles were picked using DogPicker^64^ and 3D classification was done using Relion 3.0^65^.

### Visualization and statistical analysis

Data visualization and statistical analysis were performed in GraphPad Prism Software (version 8.3).

