## Supplementary Materials for "Broad SARS-CoV-2 Neutralization by Monoclonal and Bispecific Antibodies Derived from a Gamma-infected Individual"

**This PDF file includes:**

Supplementary Figures 1 and 2

Supplementary Tables 2 and 3

**Other Supplementary Information for this manuscript include the following:**

Supplementary Table 1 (separate excel file)

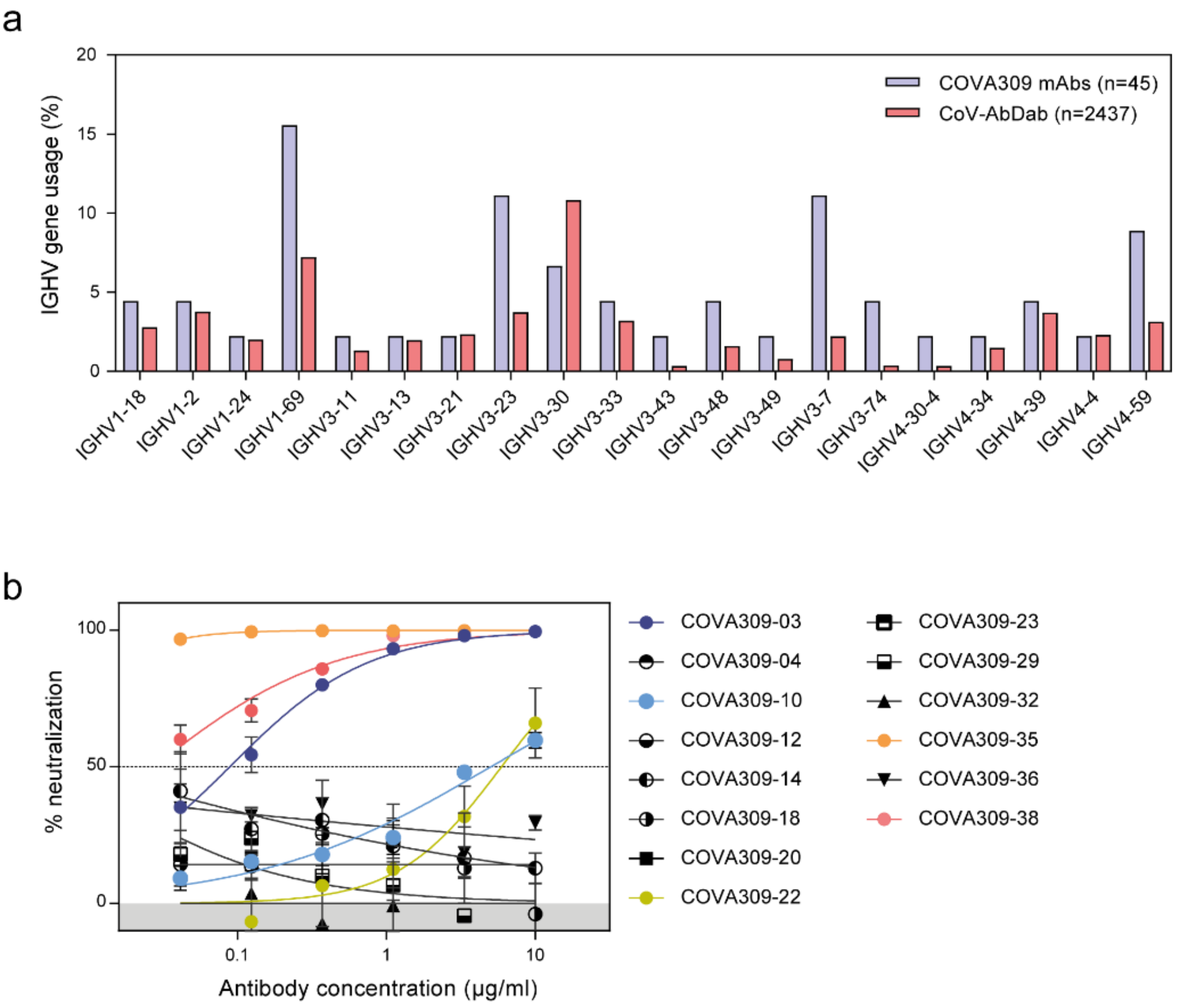


**Supplementary Figure 1. a.** Bar graph showing the mean IGHV gene usage (%) by the 293T-produced COVA309 mAbs (purple, n=45) versus sequences derived from WT-elicited B cells included in the CoV-AbDab database[30] (red, n=2437). **b.** Percentage of neutralization of SARS-CoV-2 Gamma pseudovirus by the 14 HEK293F-produced COVA309 mAbs. COVA309-03, -10, -22, -35 and -38 showed considerable neutralizing activity against the autologous variant and were selected for further analysis.

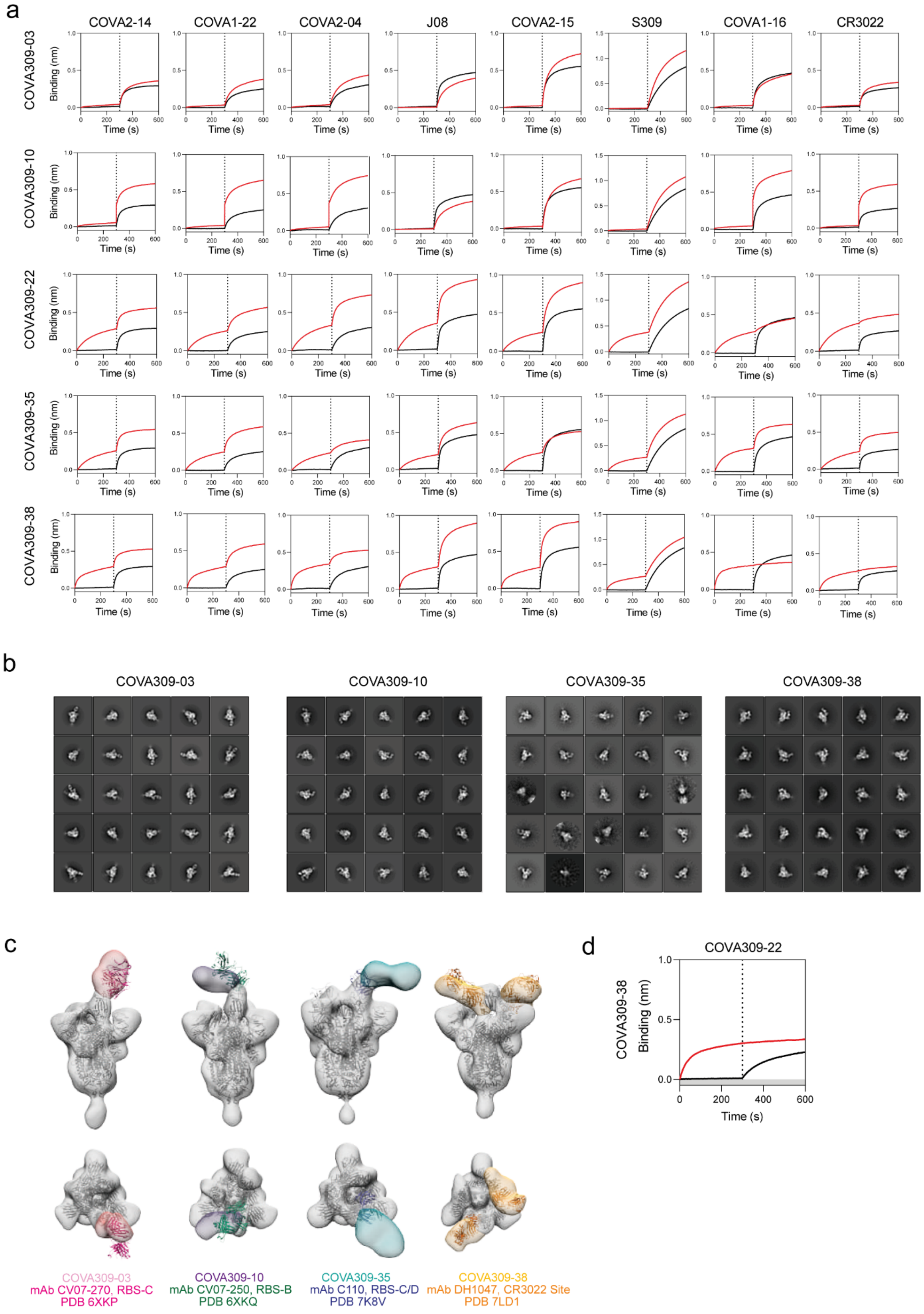
a
COVA2-14
COVA1-22
COVA2-04
J08
COVA2-15
S309
COVA1-16
CR3022
COVA309-03
COVA309-10
COVA309-22
COVA309-35
COVA309-38
Binding (nm)
Time (s)
b
COVA309-03
COVA309-10
COVA309-35
COVA309-38
c
COVA309-03
mAb CV07-270, RBS-C
PDB 6XKP
COVA309-10
mAb CV07-250, RBS-B
PDB 6XKQ
COVA309-35
mAb C110, RBS-C/D
PDB 7K8V
COVA309-38
mAb DH1047, CR3022 Site
PDB 7LD1
d
COVA309-22
COVA309-38
Binding (nm)
Time (s)

**Supplementary Figure 2. a.** COVA309 mAbs competition to known SARS-CoV-2 mAbs, including COVA2-14 (S2), COVA1-22 (NTD), COVA2-04 (RBS-A), J08 (RBS-B), COVA2-15 (RBS-D), S309 (S309 site), COVA1-16 (CR3022 site) and CR3022 (CR3022 site)[6,41-43], as assessed by BLI. Black curves represent the baseline binding of the competitors, while the red curves show the binding of the competitors in the presence of the analytes. **b.** Representative 2D images from NS-EM analysis of Omicron 6P and Gamma 6P S trimers complexed with the COVA309-03, -10, -35 and -38 Fabs. **c.** Reference mAb models fitted into COVA309 mAbs density for RBS class assignment. mAbs C110 (blue; PDB:7K8V), CV07-250 (green; PDB: 6XKQ), CV07-270 (pink; PDB: 6XKP), DH1047 (orange; PDB: 7LD1)[44-45]. **d.** Cross-competition of COVA309-38 and COVA309-22 for binding to SARS-CoV-2 WT S, as measured by BLI assay. Red curve indicates competition between the mAbs, while the black curve corresponds to COVA309-22 S binding in the absence of the analyte.

### Supplementary Table 2. X-ray data collection and refinement statistics

| **Data collection** | COVA309-22 + SARS-CoV-2 RBD |
|---|---|
| Beamline | SSRL12-1 |
| Wavelength (Å) | 0.9795 |
| Space group | $P\ 2_1\ 2_1\ 2_1$ |
| Unit cell parameters | |
| a, b, c (Å) | 54.7, 152.5, 240.8 |
| α, β, γ (°) | 90, 90, 90 |
| Resolution (Å) [a] | 50.0-3.70 (3.76-3.70) |
| Unique reflections [a] | 22,407 (2,111) |
| Redundancy [a] | 12.4 (9.5) |
| Completeness (%) [a] | 99.8 (98.9) |
| $<I/\sigma_I>$ [a] | 4.9 (0.9) |
| $R_{sym}$[b] (%) [a] | 37.6 (>100) |
| $R_{pim}$[b] (%) [a] | 11.1 (57.7) |
| $CC_{1/2}$[c] (%) [a] | 99.6 (81.3) |
| **Refinement statistics** | |
| Resolution (Å) | 46.8-3.70 |
| Reflections (work) | 22,382 |
| Reflections (test) | 2,111 |
| $R_{cryst}$[d] / $R_{free}$[e] (%) | 22.6/27.5 |
| No. of copies in ASU | 3 |
| No. of atoms | 14,610 |
| Fab | 9,939 |
| RBD | 4,671 |
| Average *B*-values ($Å^2$) | 81 |
| Fab | 82 |
| RBD | 81 |
| Wilson *B*-value ($Å^2$) | 87 |
| **RMSD from ideal geometry** | |
| Bond length (Å) | 0.002 |
| Bond angle (°) | 0.57 |
| **Ramachandran statistics (%)** | |
| Favored | 92.2 |
| Outliers | 0.06 |
| **PDB code** | pending |

[a] Numbers in parentheses refer to the highest resolution shell.

[b] $R_{sym} = \Sigma_{hkl} \Sigma_i | I_{hkl,i} - <I_{hkl}> | / \Sigma_{hkl} \Sigma_i I_{hkl,i}$ and $R_{pim} = \Sigma_{hkl} (1/(n-1))^{1/2} \Sigma_i | I_{hkl,i} - <I_{hkl}> | / \Sigma_{hkl} \Sigma_i I_{hkl,i}$, where $I_{hkl,i}$ is the scaled intensity of the $i^{th}$ measurement of reflection h, k, l, $<I_{hkl}>$ is the average intensity for that reflection, and *n* is the redundancy.

[c] $CC_{1/2}$ = Pearson correlation coefficient between two random half datasets.

[d] $R_{cryst} = \Sigma_{hkl} | F_o - F_c | / \Sigma_{hkl} | F_o | \times 100$, where $F_o$ and $F_c$ are the observed and calculated structure factors, respectively.

[e] $R_{free}$ was calculated as for $R_{cryst}$, but on a test set comprising 5% of the data excluded from refinement.

[f] From MolProbity[66].

**Supplementary Table 3. Mutations present in the S protein constructs and pseudoviruses compared to the WT strain.**

| Alpha | Beta | Gamma | Delta | Omicron BA.1 | Omicron BA.2 |
|---|---|---|---|---|---|
| deletion (Δ) of H69-V70<br>ΔY144<br>N501Y<br>A570D<br>D614G<br>P681H<br>T716I<br>S982A<br>D1118H | L18F<br>D80A<br>D215G<br>L242H<br>R246I<br>K417N<br>E484K<br>N501Y<br>D614G<br>A701V | L18F<br>T20N<br>P26S<br>D138Y<br>R190S<br>K417T<br>E484K<br>N501Y<br>D614G<br>H655Y<br>T1027I | T19R<br>G142D<br>E156G<br>Δ157-158<br>L452R<br>T478K<br>D614G<br>P681R<br>D950N | A67V<br>Δ69-70<br>T95I<br>G142D<br>Δ143-145<br>Δ211<br>L212I<br>ins214EPE<br>G339D<br>S371L<br>S373P<br>S375F<br>K417N<br>N440K<br>G446S<br>S477N<br>T478K<br>E484A<br>Q493K<br>G496S<br>Q498R<br>N501Y<br>Y505H<br>T547K<br>D614G<br>H655Y<br>N679K<br>P681H<br>N764K<br>D796Y<br>N856K<br>Q954H<br>N969K<br>L981F | T19I<br>Δ24-26<br>A27S<br>G142D<br>V213G<br>G339D<br>S371F<br>S373P<br>S375F<br>T376A<br>D405N<br>R408S<br>K417N<br>N440K<br>S477N<br>T478K<br>E484A<br>Q493R<br>Q498R<br>N501Y<br>Y505H<br>D614G<br>H655Y<br>N679K<br>P681H<br>N764K<br>D796Y<br>Q954H<br>N969K |
